# Needle in a haystack: A droplet digital polymerase chain reaction assay to detect rare helminth parasites infecting natural host populations

**DOI:** 10.1101/2025.01.23.634533

**Authors:** Chloe A. Fouilloux, Eric Neeno-Eckwall, Ipsita Srinivas, Jonathan Compton, Josh Sampson, Jesse Weber, Cole Wolf, Amanda Hund, John Berini, Heather Alexander, Emma Choi, Daniel I. Bolnick, Jessica L. Hite

## Abstract

Helminths infect humans, livestock, and wildlife, yet remain understudied despite their significant impact on public health and agriculture. Because many of the most prevalent helminth-borne diseases are zoonotic, the health of diverse host species are closely interconnected. Therefore, understanding helminth transmission among wildlife could improve predictions and management of infection risks across species. A key challenge to understanding helminth transmission dynamics in wildlife is accurately and quantitatively tracking infection levels across hosts and environments. Traditional methods, such as visual parasite identification from environmental samples or infected hosts, are time-consuming, while standard molecular techniques (e.g., PCR and qPCR) often lack the sensitivity to reliably detect lower parasit burdens. These limitations often underestimate the prevalence and severity of infection, hindering efforts to manage infectious diseases. Here, we developed a multiplexed droplet digital PCR (ddPCR) assay to quantify helminth levels in aquatic habitats using 18S rRNA target genes. Using *Schistocephalus solidus* and their copepod hosts as a case study, we demonstrate ddPCR’s sensitivity and precision. By establishing a 1:1 infection standard in the lab, we contextualize ddPCR gene concentration data to quantify both host and parasite numbers in field samples. The assay is highly reproducible, reliably detecting target genes at concentrations as low as 1 picogram of DNA in lab standards and field samples (multi-species and eDNA). Thus, we provide a toolkit for quantifying infection loads in intermediate hosts and monitoring infection dynamics across spatio-temporal scales in multiple helminth systems of concern for public health, agriculture, and conservation biology.

**Graphical abstract:** Applications of ddPCR probe-primer design to parallel systems.
Cyclopoid copepods serve as initial hosts for diverse helminthic diseases distributed globally. The primers designed in this assay are suitable for other systems, with minimal work required for probe design specific to each helminth species.

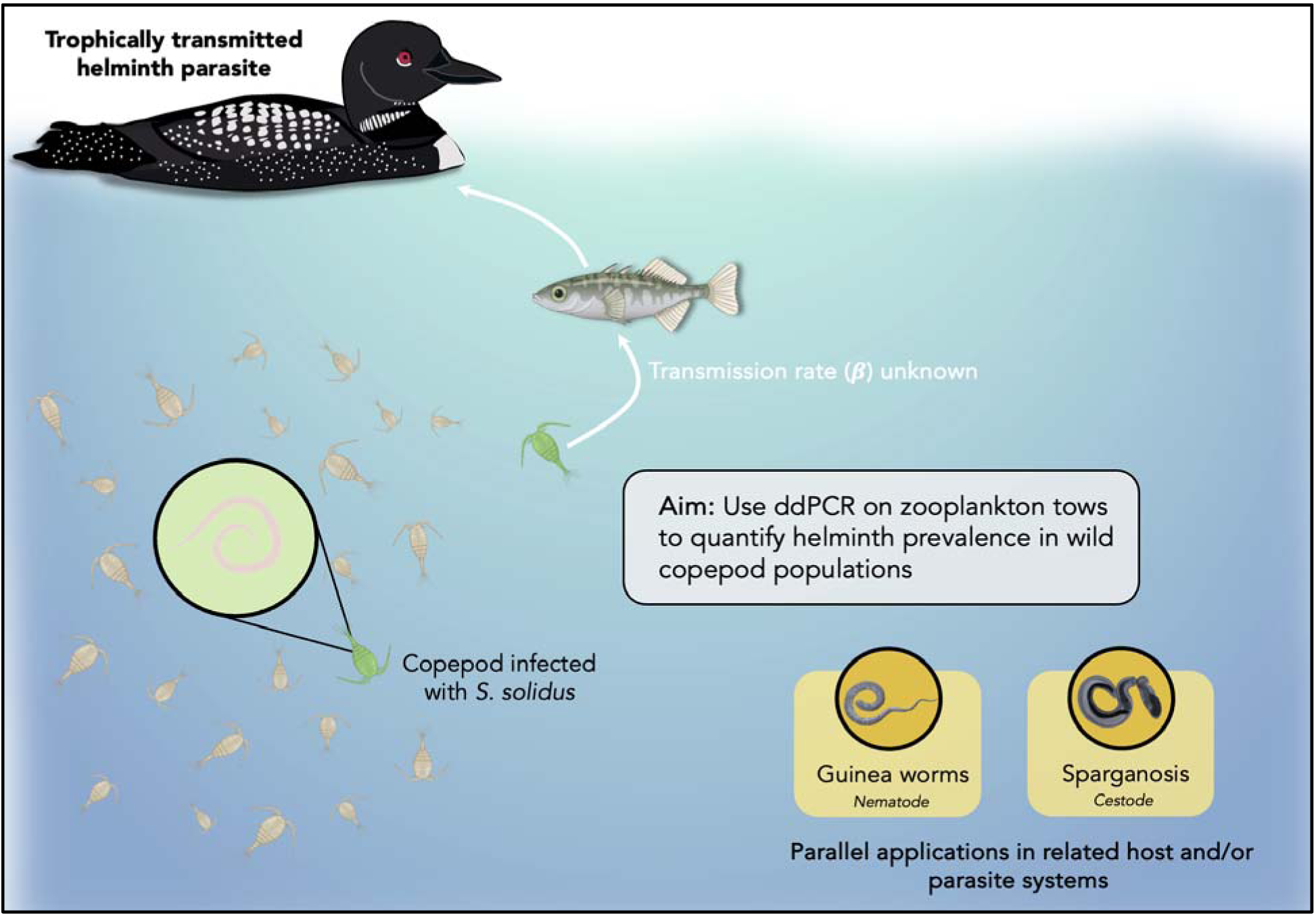

## Introduction

From populations to landscapes, parasites can drastically alter the ecology of an ecosystem. Parasites with complex life cycles can have particularly drastic consequences for global health, as they infect hosts that span multiple trophic levels (Labaude et al. 2015, Barber et al. 2016). Utilizing diverse hosts across their life cycle makes these parasites difficult to track, and considerable work has been invested to disentangle the transmission dynamics of parasites in multi-host systems (Fenton et al. 2015, Webster et al. 2016). Yet, a black box often exists around infection in first-intermediate hosts in many host-parasite systems, due in large part to the difficulty of detection and identification of microscopic larval parasites (Kurtz et al. 2002, Fenton et al. 2015, Bass et al. 2021, Klawonn et al. 2023). While a parasite’s downstream effects can be quantified in second-intermediate or definitive hosts (when parasites are mature and macroscopic), the evolutionary forces shaping initial infection and the ecology maintaining endemic infection levels (Benesh 2016) remain difficult to assess on a population level.

Helminth parasites (roundworms, tapeworms, and flukes) remain understudied despite the enormous challenges they pose to global health and livestock sectors (Lustigman et al. 2012, Charlier et al. 2014), partially because many helminth life cycles begin in invertebrate hosts that are challenging to study and control (Scholz et al. 2009). As a consequence of economic losses and compromised human and animal welfare, there have been repeated calls to improve detection of early infection and to quantify helminths in dynamic, ecologically-realistic environments (Lustigman et al. 2012, Mbong Ngwese et al. 2020). However, helminths remain challenging to study (and manage) for multiple interrelated reasons.

For example, *Diphyllobothrium* has infected humans for millennia (its oldest identified human host was a mummified corpse from ancient Chile; Reinhard and Urban 2003). Despite this long history of human infection, the multihost life cycle of the parasite was only described in the 20th century, when copepods (an aquatic crustacean) were shown to be their intermediate host (Scholz et al. 2009). An especially debilitating helminth parasite, *Dracunculus medinensis,* infected over 3.5 million humans in the 1980s (Cairncross et al. 2002). In spite of extensive control efforts, this tapeworm persists in animal reservoirs, thanks in part to the inability to monitor the initial zooplankton host (Goodwin et al. 2022). In these and other parasites with complex, multi-host lifecycles, monitoring infection levels and predicting patterns continues to perplex theoreticians and epidemiologists alike (Fenton et al. 2015, Walker et al. 2017). A lack of resolved data across sequential hosts hinders both model verification and refinement and thus, effective parasite management. Consequent economic losses and compromised human and animal welfare have inspired a call to improve detection of early infections and quantify helminths in dynamic, ecologically realistic environments (Lustigman et al. 2012, Mbong Ngwese et al. 2020).

Disentangling infection dynamics hinges on the ability to track host populations and their corresponding infection levels. Assessing these relationships in helminthiases remains challenging, because, unlike many other parasites, helminths are difficult to culture and study under standard laboratory conditions (Brindley et al. 2009). Consequently, most of our knowledge surrounding helminth infections is based on patterns derived from the analysis of field samples. Conventional diagnostic methods primarily depend on microscopy for parasite identification. While arguably more affordable, microscopy is both labor intensive and ineffective in reliably detecting early stages of infection within hosts and early life history stages of parasites (Boonham et al. 2020, Mbong Ngwese et al. 2020).

Molecular approaches such as qPCR have improved the specificity and sensitivity of diagnostic assays; however, they do not provide absolute quantification of targets, are prone to inhibition by environmental contamination (Shannon et al. 2007), and are unreliable when detecting minute amounts of DNA (Amoah et al. 2017). Additionally, critics of qPCR highlight that problems with data reproducibility underscore the need for new and improved quantitative methods (Hindson et al. 2011, Dijkstra et al. 2014, Bustin et al. 2013). A lack of quantification hinders disease containment efforts and restricts the empirical data required to predict environmental drivers associated with increased infections in nonhuman hosts (Boonham et al. 2020, Goodwin et al. 2022).

Here, we use a model host-parasite system as a case study to develop and refine a sensitive molecular assay to detect and quantify infection levels in wildlife hosts and across natural environments. Specifically, we used droplet digital PCR (ddPCR) technology to design a multiplexed assay based on universal 18s rRNA primers. A multiplexed ddPCR enables rapid, high-throughput quantification of two distinct targets, allowing precise measurement of gene concentrations in both hosts and parasites. ddPCR has multiple advantages over other quantification methods. First, it provides a direct and independent quantification of DNA without standard curves (Hou et al. 2023). This approach is especially powerful for investigating newly emerging and relatively understudied parasites, when culturing and creating a known concentration of target DNA may be logistically impossible. Second, by sample partitioning and endpoint detection, ddPCR analytics quantify nucleic acids independent of reaction efficiency (Taylor et al. 2017). Free of this limitation, accurate detection can occur at much lower DNA concentrations in samples that do not require dilution to exclude possible contaminants. Third, the high sensitivity of ddPCR can capture target DNA spanning a wide gradient, especially at lower concentrations (Hiillos et al. 2021).

Despite being commercially available for over a decade (Hindson et al. 2011), the application of ddPCR remains primarily utilized in medical research. Meanwhile, the fields of disease ecology and epidemiology have called for a revamping of quantitative methods (Momčilovič et al. 2019, Boonham et al. 2020), where fundamental disease questions, such as “when” and “where” disease outbreaks occur demand sensitive and precise detection methods. Indeed, providing a solid quantitative toolkit for tracking infection levels across diverse hosts and heterogeneous environments could greatly bolster management efforts of Neglected Tropical Diseases (NTDs) globally (Brindley et al. 2009). As detecting infection levels across complex field samples is akin to finding “a needle in a haystack,” ddPCR provides a powerful solution to detecting rare DNA sequences in multi-species and environmental samples.

In this study, we empirically apply a ddPCR approach to field samples where infection levels in the initial hosts were unknown and extend the application of our designed primers to related helminthiases of global health concern. To refine this method and demonstrate its utility, we use a model host-parasite system with abundant genetic and ecological resources: *Schistocephalus solidus*, a trophically transmitted helminth, and one of its first-intermediate hosts, the cyclopoid copepod *Acanthocyclops robustus*. This system serves as a case study for disease ecology and public health. *Schistocephalus solidus* was the first parasite with a complex life cycle that was described (Abildgaard 1790) and has since emerged as a powerful model for studying host-parasite co-evolution (Barber and Scharsack 2009) and conserved immune responses like fibrosis (Hund et al. 2022).

More broadly, the *S. solidus*-copepod system shares similarities with other neglected tropical diseases (NTDs). Like other helminths (e.g., Guinea worms, Box 1), *S. solidus’* first intermediate host is also a cyclopoid copepod. As both an (ecto-)parasite and host to hundreds of parasite species, copepods profoundly affect wildlife, aquaculture, and humans (Bass et al. 2021). However, most knowledge of parasite dynamics is limited to vertebrates, leaving critical gaps in understanding invertebrate population dynamics, infection risk to secondary and tertiary hosts, and overall epidemiological dynamics. To address these gaps, we outline applications of our ddPCR toolkit for other multi-host parasites, including relatives of *S. solidus* (family Diphyllobothriidae) that infect humans via farmed fish (Scholz et al. 2019). Together, our study highlights ddPCR as a uniquely reliable quantitative tool, offering new insights across the field of disease ecology.

**Box 1.**
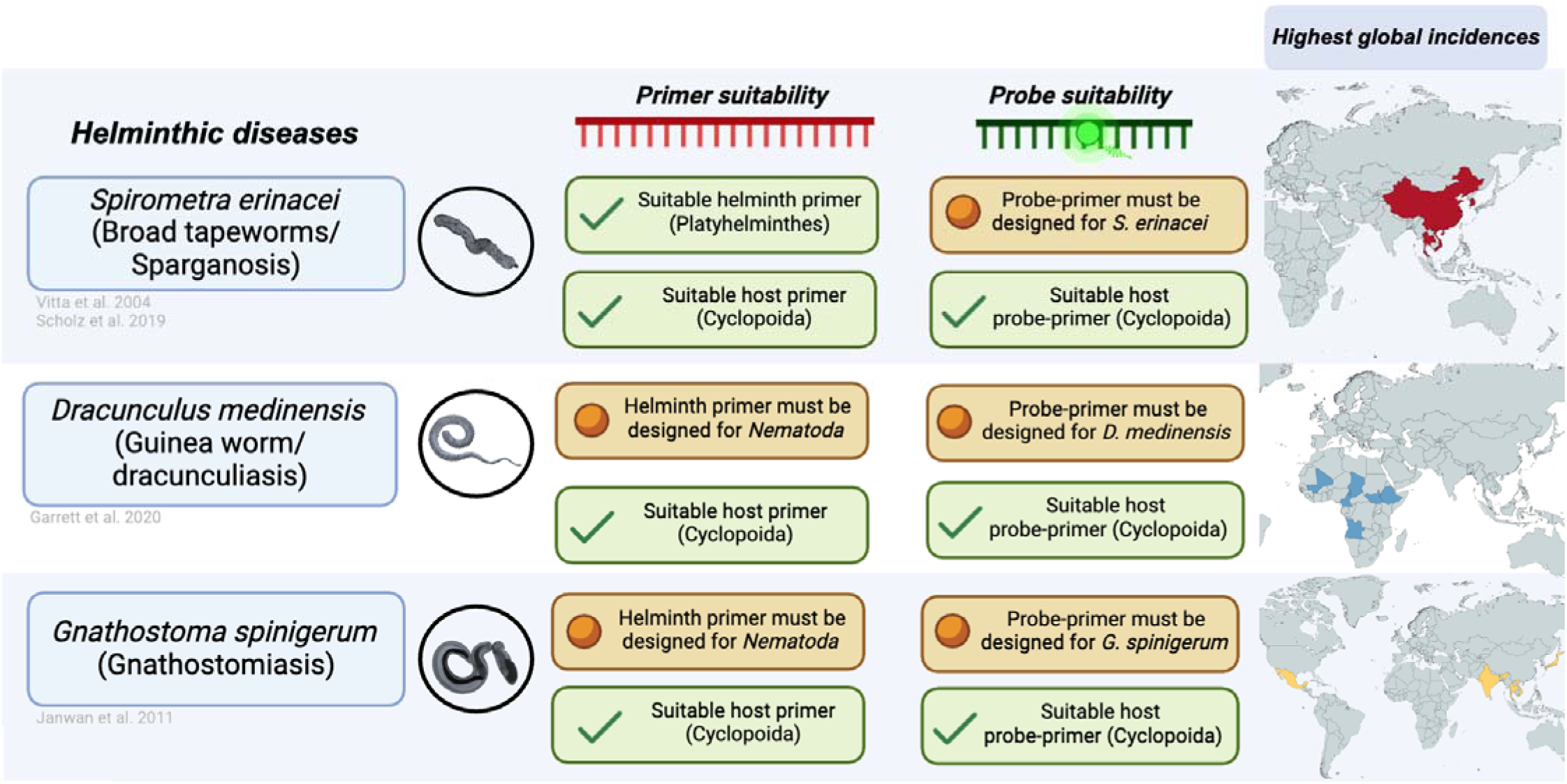
Applications of ddPCR probe-primer design to parallel systems. Cyclopoid copepods serve as initial hosts for diverse helminthic diseases distributed globally. The primers designed in this assay are suitable for other systems, with minimal work required for probe design specific to each helminth species.

## Materials and Methods

### Natural history of the focal host-parasite system

Like many other helminths, *S. solidus* is characterized by a complex life cycle requiring transmission between more than one host species (Wedekind 1997). Tapeworm eggs hatch from freshwater substrate, yielding free-swimming larvae (coracidia) that are consumed by and infect multiple species of cyclopoid copepods (Wedekind 1997). This parasite develops into procercoids within copepods and become capable of infecting its secondary obligate host, the threespine stickleback fish (*Gasterosteus aculeatus*) (Nishimura et al. 2011). The final stage of the parasite’s life cycle is reached when infected stickleback are consumed by piscivorous birds (Wedekind 1997).

### Field sites and field sample collections

We sampled three lakes from June-August 2023 across Vancouver Island, B.C., Canada: Pachena Lake (GPS = 48.834893, -125.03362; 54.9 ha), Black Lake (GPS = 48.761545, - 125.101215; 69.5 ha), and Blackwater Lake (GPS= 50.1684, -125.5916; 37.5 ha). These lakes were selected because they spanned a gradient in lake-level mean parasite loads of *S. solidus’* obligate second host, the threespine stickleback (<u>Supplementary File C</u>). We visited lakes monthly and at each visit, we collected zooplankton samples and eDNA. We collected zooplankton samples with vertical tows from the epilimnion (i.e., the upper, warmer layer of stratified lakes; Wetzel 2001). We used an EXO2 multiprobe sonde (YSI Incorporated, Ohio USA) to identify the epilimnion. We pooled three vertical tows of a Wisconsin net (13 cm diameter, 80-μm mesh). Samples were immediately stored in 95% ethanol and transported to University of Wisconsin, Madison where they were stored at room temperature for two months and then moved to -4°C for long-term storage. During each sampling visit, and each location used for the zooplankton tows in the epilimnion, we also collected eDNA samples. We vacuum filtered 4 liters of lake water through a self-preserving 5-μM filter (filter: Smith-Root 11580-25, Pump: Smith-Root 12099; Thomas et al. 2018). Filters were stored at room temperature and transported to the University of Connecticut.

### Experimental infection assays

To translate 18S gene copy numbers quantified by ddPCR into actual animal and parasite numbers, we created a 1:1 infection standard using controlled infection assays. Standards were composed of 100 individual adult *Acanthocyclops robustus* (Cyclopidae) copepods confirmed to be infected with a single *S. solidus* coracidium (n = 100) from exposure seven days prior. In parallel, we also ran a “control” standard of 100 unexposed adult copepods to ensure that multiplexing ddPCR reactions did not impact the quantification of copepod genes.

All hosts used for the infection assay were selected from existing cultures isolated from Echo Lake (Vancouver Island, B.C., Canada) in 2015/2016. Prior to beginning the infection assays, laboratory cultures of copepod hosts were maintained in 1 L flasks at 19L with a 16:8 L:D cycle with 900 mL of standard (low-hardness) COMBO water for animals (artificial lake water media, Kilham et al. 1998). Animal stocks were fed freeze-dried crushed *Artemia* (AMZEY Natural Artemia; 0.022 mg L^-1^) every other day.

Parasite eggs were originally collected from Kjerag Fjord, Norway (8 September 2022, GPS: 67.501487, 14.742647). To minimize fungal growth, eggs were washed several times with sterile water (Weber et al. 2017) and maintained in long-term storage in the darkness at 4°C in 15mL falcon tubes at the University of Wisconsin, Madison. To stimulate egg hatching, 200µl of *the* egg suspension was aliquoted into a single well of a foil-covered 24-well microtiter plate with 2mL of COMBO media and incubated in the dark at 18°C for seven days. Following this, egg plates were moved to room temperature (25-26°C) and placed under full-spectrum grow lights (GT-Lite LED Grow Bulb; 13.09 PPF, 8.5 Watts; SKU:GR-A19) with a 16:8 L:D cycle (Jakobsen et. al, 2012; Weber et al., 2017).

On the day prior to parasite exposure, single adult copepod hosts were isolated individually and maintained in the 24-well plates with 1.5mL COMBO at 19□ under a 16:8 L:D cycle (Jakobsen et al. 2012). To improve infection success rates, which relies on hosts ingesting parasite eggs, hosts were maintained without food 24-hours before exposure. Following starvation, one coracidium was placed in each well for a 24-hour exposure period. Immediately following the 24-hour exposure, isolated individuals were fed aliquots of 1 mL of *Artemia* suspension from a 0.022 mg L^-1^ stock solution every other day. Single infections were confirmed seven days post-exposure using a compound microscope (Leica M80 at 60x magnification using a Leica KL 300 LED). Infected hosts were placed individually into a 1.5 mL Eppendorf tube and stored at -20°C until DNA extraction.

### DNA Extraction

DNA from zooplankton was extracted using PowerFecal Pro Kits (Qiagen 51804) following manufacturer protocol, with an additional 10-minute 65□ heating step to help lyse zooplankton. DNA quantity and quality were measured using a NanoDrop (ND-1000) Spectrophotometer and extracts were stored at -20□. DNA used for this experiment came from 10ng/µl aliquots. eDNA from water filters was extracted using the DNAeasy PowerWater kit (QIAGEN, United States; see Kraemer et al. 2020 for details). DNA extracted from water filters was visually inspected for intactness using agarose gels and quantified using a Qubit assay (Lumiprobe, United States).

### Design of 18S primers for both Schistocephalus solidus and copepods

While *S. solidus* obligately infects threespine stickleback fish (Bråten 1966), there are a range of cyclopoid copepods that may eat larval stages of the parasite (Barber and Scharsack 2009). We designed primers for copepods that were broad enough to amplify diverse Cyclopoids but excluded other zooplankton (Table 1). We tested primers that have been previously reported to amplify copepod sequences (Hubbard et al. 2016, Teterina et al. 2016, Mercado-Salas et al. 2021) but none either (a) consistently amplified copepod-rich samples from Vancouver Island lakes or (b) were the correct size and melting temperature required for ddPCR.

**Table 1.**
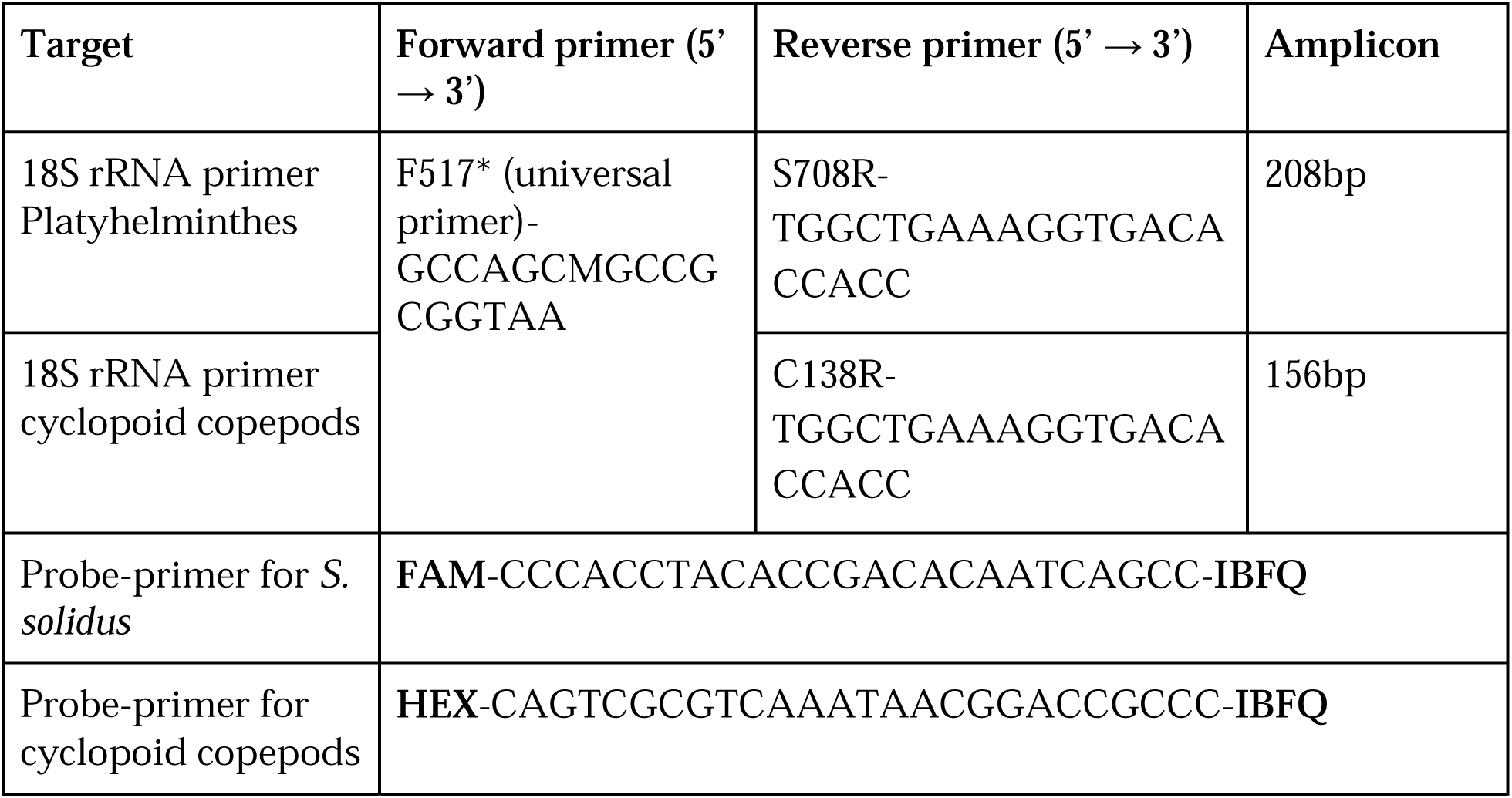
Primers and probe primers optimized for ddPCR assay. Bolded values denote the relative positions of the fluorophore and quencher used in ddPCR probe-primers.

18S rRNA genes were chosen because they contain conserved sequences surrounding variable regions that are species-specific; for both *S. solidus* and copepods we use the universal 18S primer, F517 as the forward sequence (Bates et al. 2012, Table 1). We designed primers for *S. solidus* using a complete 18S rRNA sequence available on NCBI (*GenBank: AF124460.1).* Having accessed this sequence, we aligned universal 18S primers to *S. solidus* (F517, R1119, Bates et al. 2012). We performed a local BLAST on the resulting amplicon against two assemblies of *S. solidus* available in GenBank (GCA_900618435.1, GCA_017591395.1) to identify 18S rRNA present in assemblies. The resulting hits were aligned using AliView (v. 1.28, Larsson 2014). From these aligned assemblies, we visually identified a 21bp conserved region common among platyhelminthes that serves as a reverse primer (S708R; 5’- TGGCTGAAAGGTGACACCACC-3’). This primer was designed to be broad, as the probe-primer of the ddPCR provides an additional level of specificity to bind only closely related *Schistocephalus* species (<u>Supplementary File</u> A).

Copepod primers were similarly designed using the same 18S universal forward primers used for *S. solidus* (Table 1). Using the 18S rRNA sequence available for the cyclopoid copepods, *Acanthocyclops viridis* (GenBank: AY626999.1), we identified the amplified region, and BLASTed the amplicon against the nucleotide collection (nr/nt database) in NCBI. From selected alignments, we found a region where universal primers would bind to all aligned copepod sequences. We aligned the first 101 hits from this search and identified a conserved region to serve as the reverse primer (C138R, 5’-TGGCTGAAAGGTGACACCACC-3’). The validity of primers were assessed using primer BLAST on NCBI. PCR copepod primers will amplify diverse genera of copepods though more strongly amplify cyclopoids (see <u>Supplementary File</u> B).

PCR assays using 18S primers were conducted on genomic DNA samples to ensure primer specificity. PCR reactions were performed in a total volume of 25μL [12.5μL Master Mix, 8μL PCR water, 1.25μL forward and reverse primer (10μmol l^-1^ conc.), 2μL DNA (10ngμL^-1^ conc.) (or 2μL of PCR water for NTC)] in a BioRadT100 thermal cycler. PCR conditions included a denaturation step of 5 min at 95°C, followed by annealing and elongation step of 40 cycles at 95°C for 45s, 54°C for 45s, 72°C for 60s, and a final step of 72°C for 7 min with a 1°C/sec ramp speed. PCR produced were run on 1.5% agarose gel using SYBR safe staining (Invitrogen). After gel visualization, the appropriate sized bands were excised and purified from the gel using Zymoclean Gel DNA Recovery Kit (Cat No. D4001S, LOT No. 228633) following the manufacturer’s protocol. Purified PCR products were then sequenced by Functional BioSciences (Madison, WI) to ensure primers were amplifying the desired targets; sequencing data showed that these PCR products contained target segments. Primer specificity was also internally validated by using DNA extracts from lab-reared *A. robustus* (J. Weber 2023) that were experimentally exposed to larval *S. solidus*.

Sequenced *S. solidus* amplicons were identical to the genome assembly available on NCBI Gene Bank (assembly number: GCA_017591395.1). Cyclopoid primers successfully amplified both lab-reared copepods and wild copepods from mixed species zooplankton tows (where zooplankton identification in paired samples all contain cyclopoid copepods, Srinivas et al. *in prep*.).

### ddPCR assays

Our goal was to produce an assay that could be used to detect and estimate infection loads of a rare helminth parasite from total DNA extracts that contained both diverse zooplankton and target parasite DNA. ddPCR requires probe-primers to bind within the amplicon and cannot overlap either of the amplification primers. This provides an additional degree of specificity to the reaction (Hou et al. 2023).

Primer-probe suitability (e.g., melting point, GC-content, secondary structure) was checked using the OligoCalc Tool (biotools.nubic.northwestern.edu; starting settings recommended by BioRad: 300nM primer, 50mM salt, using nearest neighbor melting temperature). Here, the F517 universal primer (Bates et al. 2012) was shortened by 2nts at the 5’ end to lower its melting temperature (renamed F517). Based on the broad primers for platyhelminthes, we designed probe-primers to be specific to *S. solidus.* Experimental testing of the probe-primers demonstrates distinct amplification of *S. solidus* from locally co-occurring helminth, *S. cotti* (<u>Supplementary File</u> A). These primers do amplify the sister species *S. pungitii* which differs by a two nucleotide gap from *S. solidus;* however, these helminths have yet to be observed in the Vancouver Island area (DIB personal obs.). For copepods, we selected a 26nt probe-primer within the amplified primer region based on manual inspection of alignment. Copepods probe-primers are specific to cyclopoids and will not amplify other genera, such as calanoids (see <u>Supplementary File</u> B for empirical support). *S. solidus* probe-primers were labeled with a 5’ FAM fluorophore (5’-CCCACCTACACCGACACAATCAGCC-3’) and copepod primers were labeled 5’ HEX fluorophore (5’-CAGTCGCGTCAAATAACGGACCGCCC-3’) to allow multiplexed reactions. Probe-primers were designed and ordered from Integrated DNA Technologies (IDT), these included the 5’ fluorophores specified above and ZEN Double-Quenched Probes which contain a 3’ Iowa Black FQ (IBFQ) quencher and a proprietary internal ZEN quencher.

ddPCR was performed using Bio-Rad’s QX-200 Droplet Digital PCR System. Across the ddPCR run, we conducted both multiplex and singleplex reactions to address detection and inhibition questions. The multiplexed assay was used to detect both copepods and *S. solidus* within samples to better identify true positives (e.g., a positive signal for *S. solidus* without a signal for copepods is a false positive). To ensure that multiplexing reactions did not bias 18S quantification, we included singleplex reactions for both host (HEX) and parasite (FAM) targets. For parasites, this considered the quantification of parasite 18S genes from singleplex (FAM) and multiplex (FAM + HEX) reactions from infected copepod standards. For hosts, we compared 18S gene quantification from the same infected copepod standards (FAM + HEX) and unexposed copepod adult standards (n = 100 copepods) in a singleplex reaction (HEX). The reaction mix was prepared to a volume of 22 µl per sample. This was composed of 10µl of 2X ddPCR™ supermix for probes (BioRad) reagent mix and 1.2µl of FAM- and/or HEX-labeled primer/probe mixes (900nM primers/200nM probes, depending on single- or multiplex reaction) were aliquoted into a 96-well plate to which 8.8µl of DNA template (serially diluted from original concentration of 10ng/µl or RNase/DNase-free water for NTC) was then added. Serial dilutions and zooplankton tows were run in triplicate. Samples were mixed within wells by pipette mixing.

Once combined, 20 µl of the reaction mix and 70 µl of Droplet Generation Oil (Bio-Rad) were loaded into their appropriate wells in a single-use DG8 cartridge. Cartridges were loaded into a QX200 Droplet Generator (Bio-Rad), where samples are partitioned into nanoliter-sized droplets. 40 µl of the resulting emulsion were manually transferred to a ddPCR 96-well PCR plate (Bio-Rad), which was heat-sealed with a foil cover. The droplets were then subject to thermocycling using a Bio-Rad C1000 thermocycler with a ramp rate of 1°C/s using the following specifications: a 10 min enzyme activation step at 95°C, followed by 40 cycles of 30s at 94°C (denaturation) and 1 min at 62.5°C (annealing/extension), followed by 10 min hold at 98°C. Amplification efficiency was optimized over a temperature gradient (54.6 - 65°C), where we found the ideal optimal temperature for both primer probes at 62.5°C. All experiments included both a negative control containing nucleotide free water and a double-positive control containing *S. solidus* infected copepods. Following thermocycling, the droplets were immediately read with Bio-Rad’s Droplet Reader.

One of the advantages of ddPCR is that technical replicates are not needed as there can be more than 15,000 PCR reactions in a single well (Droplet Digital PCR Applications Guide 2017). Within a single oil droplet the presence of target DNA is assessed based on fluorescence which is ‘binned’ as either positive or negative. From this, Poisson statistics are used to estimate the absolute copy number of target DNA based on the proportion of positive droplets in the entire reaction (Jones et al. 2014). As concentration estimates hinge on droplet counts, the precision of calculations are more accurate in wells with more successfully generated/processed droplets; as such, we exclude wells with low droplet counts (<10,000 droplets) from further analysis. In the context of this study we repeated three true technical replicates in order to quantify the repeatability of detection in samples with extremely rare (< 1 picogram of total DNA) events. A threshold to separate the target positive and negative droplets is initially suggested by the ddPCR QuantaSoftware. We manually adjusted this threshold to be above the negative amplitude of both dyes (excluding more of the “rain” from Poisson calculations) for a more conservative estimate of target concentration (Fig 1); although these isolated droplets have little effect on an estimate of more than 10,000 points.

**Figure 1.**
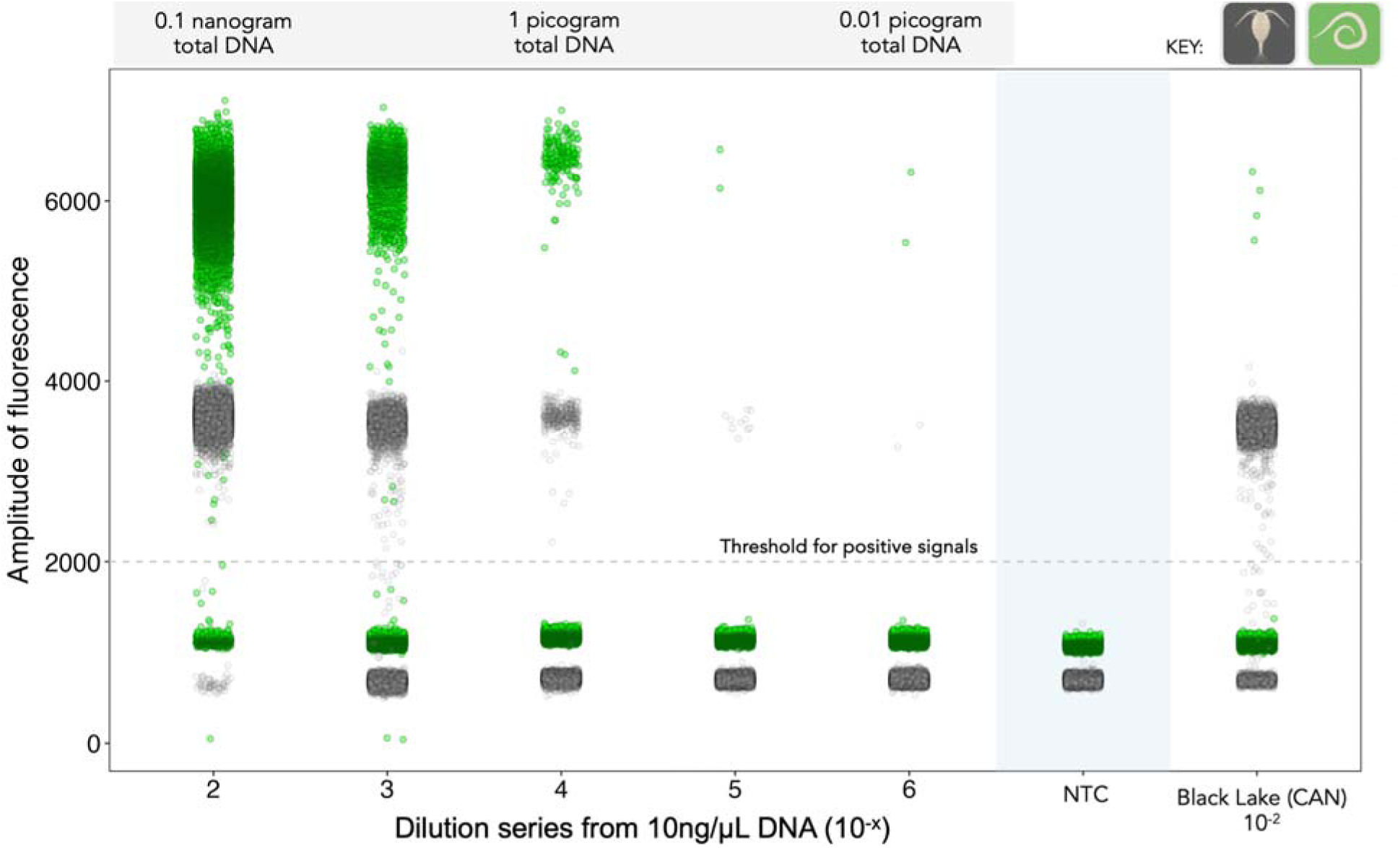
1-D plot of the limit of detection in a multiplexed ddPCR assay. Infection standard consisted of 100 singly infected copepods. Last column is a field sample included as comparison to the lab standard. The panel highlighted in blue is the negative template control. Positive reactions and negative reactions (light green and black, respectively) are separated by a manually set threshold (dashed line, at amplitude of 2000). Green points are FAM-fluorescently labeled *S. solidus* droplets and gray points are HEX-fluorescently labeled copepod droplets. Differences in fluorescence amplitude signal of each target dye (here FAM (green) shines brighter than HEX (black)) allows spatial differentiation of the droplet clusters. Each point is an individual reaction.

### Quantification of target genes

Absolute quantification of target gene copies was done with default ABS settings in QuantaSoft Analysis Pro 2.0 software (Bio--Rad). ddPCR reactions occur in droplets, which after being amplified to end-point are assigned as positive or negative for target genes, based on their fluorescence. The fraction of positive partitions within a well is used to estimate target gene concentration by modeling as a Poisson distribution which is reported in target gene copies/μL. In both 1:1 infection standard and wild zooplankton tows, undiluted DNA resulted in zero negative droplets. Without separation it is impossible to estimate target copy number. We found samples diluted to 10^-2^ (0.1 ng/μL) exhibited enough separation required for ddPCR to estimate gene concentrations.

### Limit of detection (LOD) experiment

In order to inspect the limit of detection (LOD) of this ddPCR assay, we did a 10-fold dilution series (10^-2^ to 10^-6^) from 10 ng/μl DNA stocks. Assay repeatability was determined by both (1) the % coefficient of variation (%CV = concentration standard deviation/concentration mean 100) between the replicates and (2) the linearity of the dilution assay.

## Results

### Experiment 1: What is the limit of detection of rare DNA from a known infection dose in a helminth-copepod system?

In our limit of detection experiment, we reliably (e.g., positive *S. solidus* in every replicate) detected *S. solidus* DNA in total concentrations of less than one picogram of total DNA (a 10^-5^ dilution, see Table 1 and Figure 1), otherwise expressed as a mean of 0.187 18S rRNA copies/μL. When run in triplicate, the probability of detection increases, allowing detection of true positives of *S. solidus* in as little as 0.01 picograms of DNA. The assay was reliable, demonstrating linearity across the dilution series (r^2^ = -0.98, p < 0.001, for both targets; see Figure 2). The coefficient of variation (%CV) was below 0.01 for both *S. solidus* and *A. robustus* targets. Droplet counts for the multiplexed standard experiment ranged from 12,423 to 15,834.

**Figure 2.**
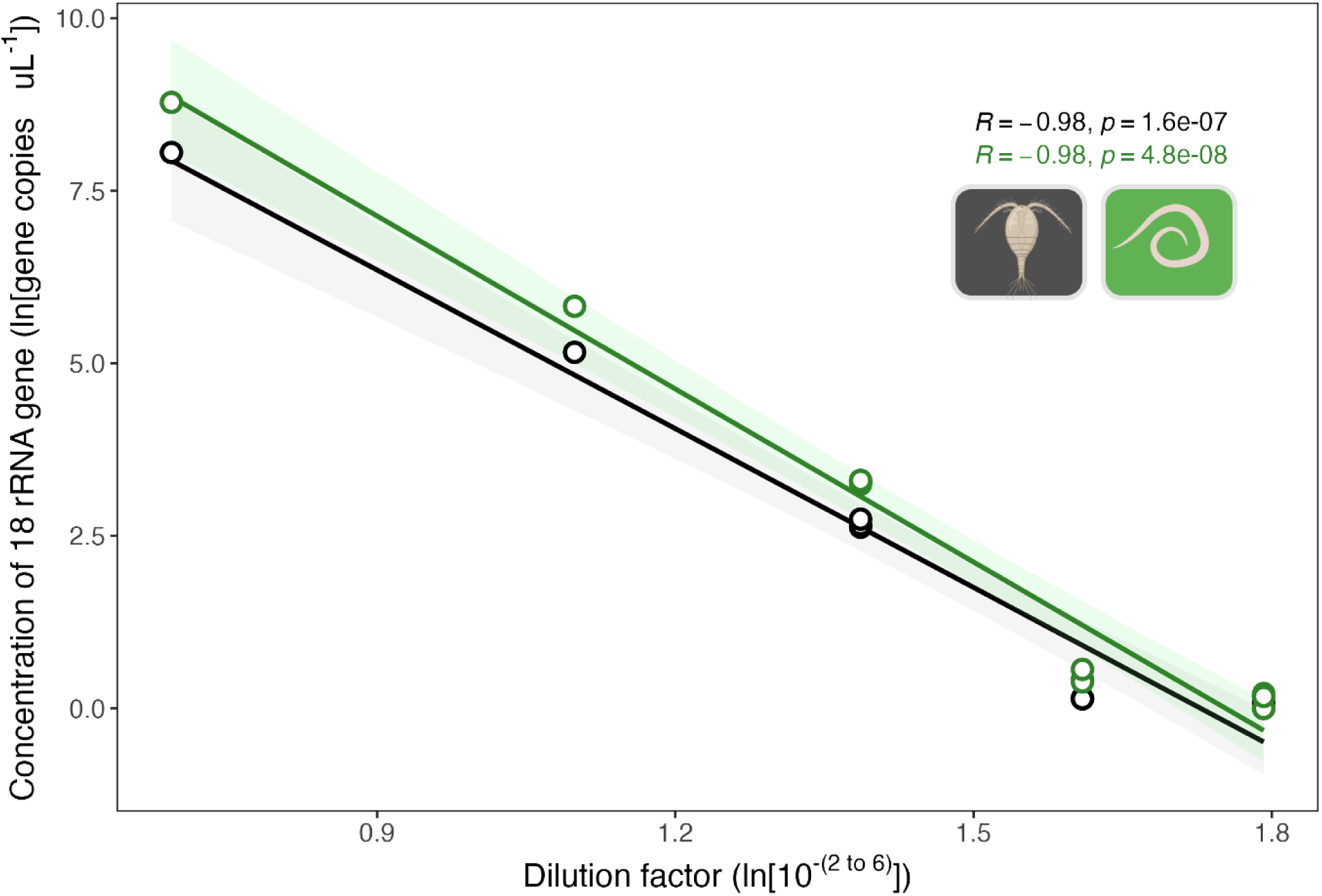
Linear relationship between dilution factor and gene concentration. There exists a significant correlation (r^2^ = -0.98, p <0.001 for both targets) between dilution and absolute concentration, demonstrating a reliable linearity of the dilution assay.

We used three independent standards to quantify zooplankton gene concentration: an infection standard that contained 100 adult *A. robustus* copepods individually infected with a single *S. solidus* and two independent standards (created from different lab lines) of 100 adult unexposed *A. robustus* copepods. We did not detect a significant difference in estimates of copepod numbers across standards (ANOVA, F-value = 1.88, p = 0.18). Indicating no effect of multiplexing or infection on copepod 18s rRNA gene quantification. When averaging copepod gene concentration of each standard at each dilution step, all estimates were on the same order of magnitude after correction (10^-4^ = 474,333 copies/μL, 10^-5^ = 330,625 copies/μL, 10^-6^ = 251,250 copies/μL). Taking the average across all three standards after correcting for dilution factor, we estimate 100 adult copepods to have 340,955 (se = 64,421) 18S rRNA gene copies/μL.

The estimate of *S. solidus* DNA was based solely on the 1:1 infection standard. Despite the %CV within each dilution step being very low, when correcting for the dilution factor we find that *S. solidus* gene estimates are an order of magnitude smaller in more dilute samples (i.e., 10^-5^ and 10^- 6^). Based on the corrected average from the less diluted replicates (10^-2^ - 10^-4^) we estimate 100 encysted *S. solidus* to have an average of 159,857 (se = 22,887) 18S rRNA gene copies/μL.

### Experiment 2: Does multiplexing ddPCR reactions inhibit the detection of rare DNA?

Using a t-test we found that there is no statistical difference between the amount of target *S. solidus* 18S DNA detected across a dilution series (0.001-0.00001ng/μL DNA) between ddPCR assays with single versus multiplexed reactions (t = 0.0759, df = 15.955, p-value = 0.940; Table 2).

**Table 2.**
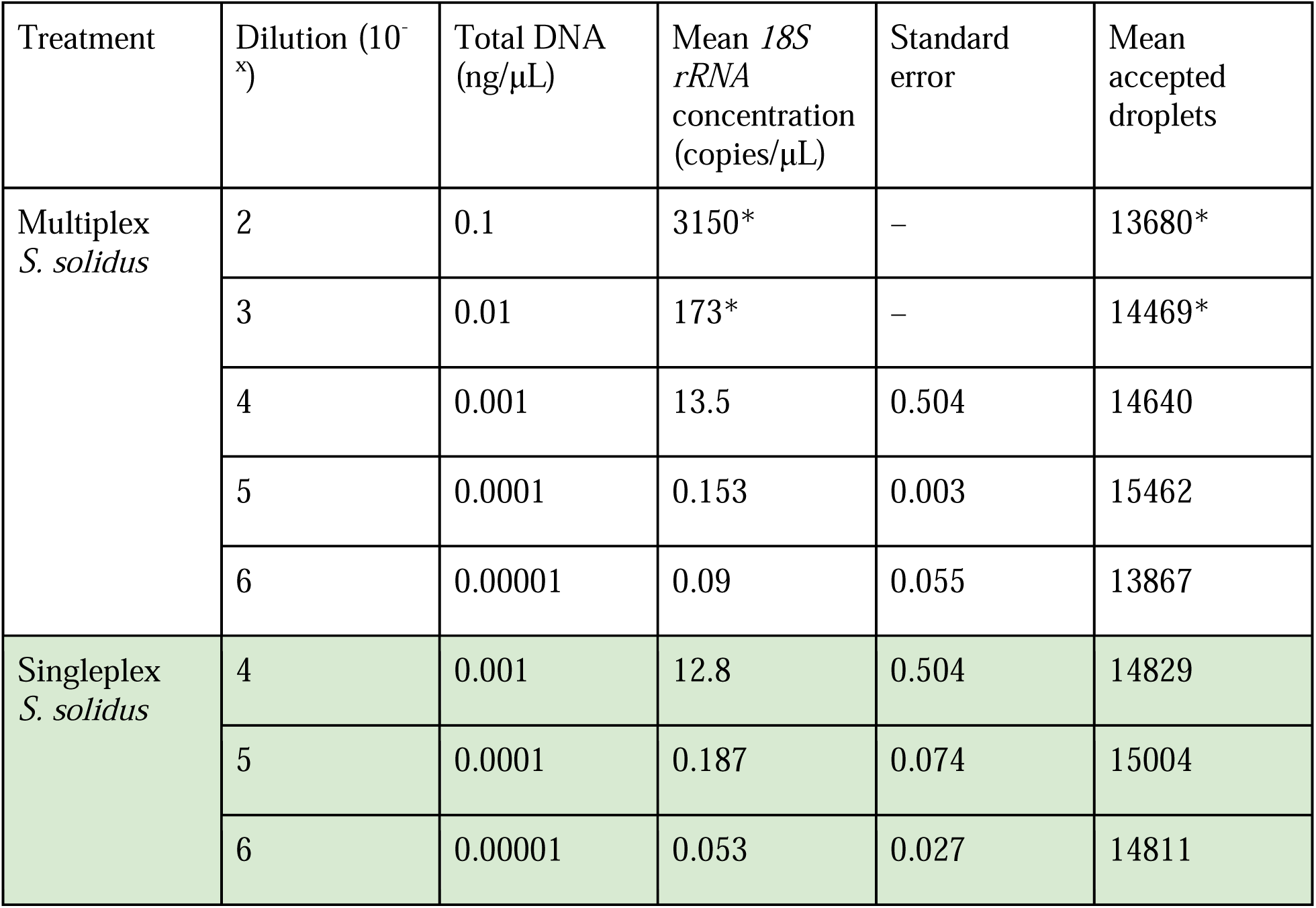
Detection of *S. solidus* DNA in single versus multiplexed reactions using an infection standard (100 copepods exposed and infected with 1 *S. solidus* parasite). Samples were diluted from 10ng/μL stock. *Indicate samples without true technical replicates.

### Experiment 3: How can we use infection standards to model infection dynamics in natural systems?

In natural systems where the mean infection load is unknown, running both 10^-1^ and 10^-2^ dilutions was ideal for both helminth detection and host quantification. We found that a 10^-1^ dilution removes potential inhibitors from a sample with minimal compromise to parasite detection. A 10^-2^ dilution generated enough separation between positive and negative droplets to quantify host density.

Using both water filters (eDNA) and zooplankton tows to ground-truth this methodology, this ddPCR assay has the sensitivity required to detect *S. solidus* within their cyclopoid copepod host from a large mixed species population (Figure 3). Using this detection system as a proof-of-concept, we calculated infection prevalence and host density across sampling In a zooplankton tow from June 2023 Pachena Lake (Vancouver Island, B.C.), we found an average of 13 copies of *S. solidus* 18S rRNA/μL (se = 0.58, corrected for dilution factor) within 108,700 copies ofμL (se = 961, corrected for dilution factor). The following spring, we found zooplankton tows from Black Lake (Vancouver Island, B.C.; sampled March 2024) contain a mean of 67.7 copies of *S. solidus* 18S rRNA/μL (se = 2.6, corrected for dilution factor) within 205,100 copies of cyclopoid 18S rRNA/μL (se = 7970, corrected for dilution factor).

**Figure 3.**
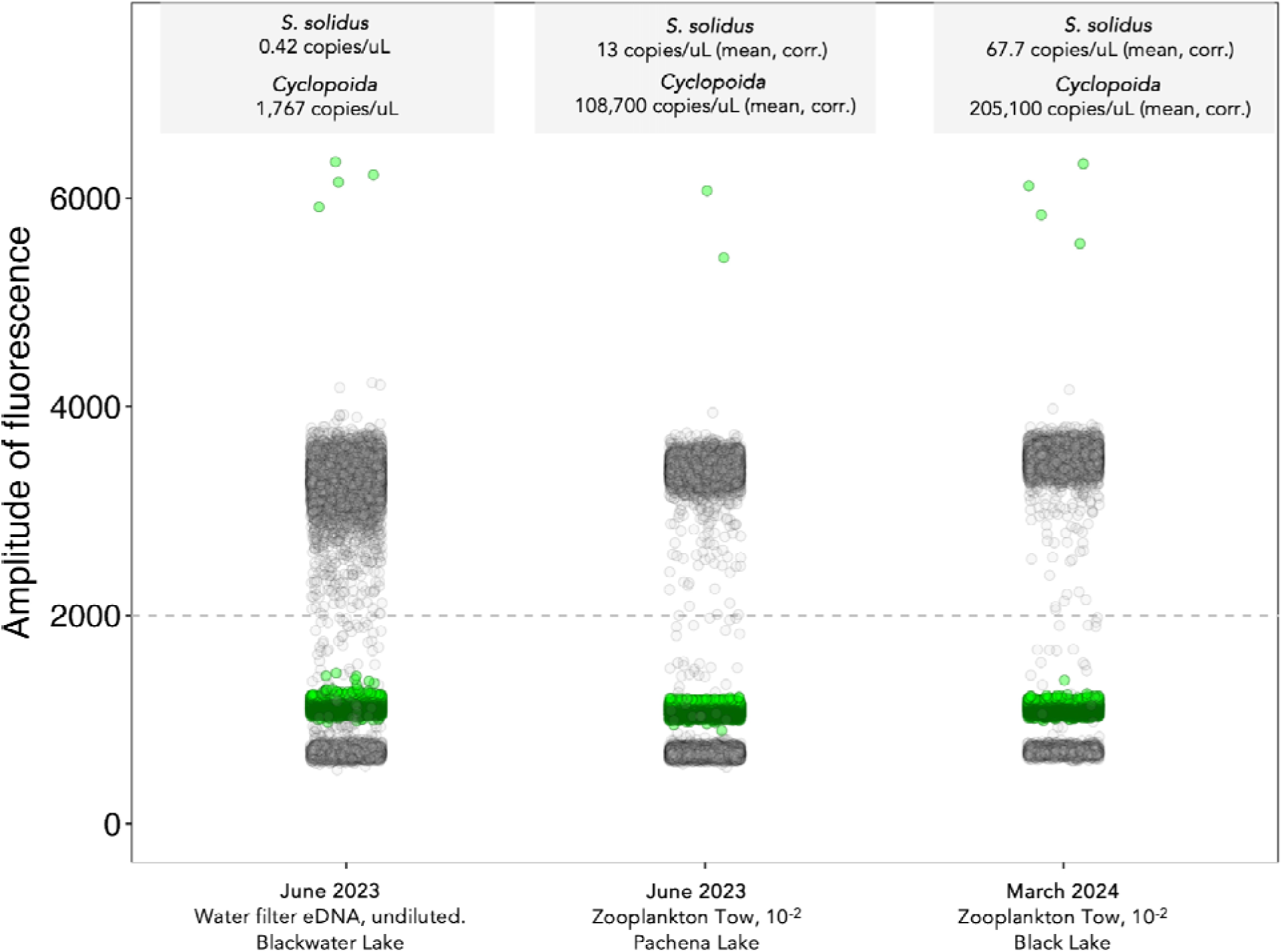
Host-parasite detection in environmental samples. In multiplexed ddPCR reactions, we find positive results in (1) diverse lakes in different months and (2) in both water samples and zooplankton tows. Single representative amplification wells are visualized here, but samples were run in triplicate across the dilution series (except for the eDNA sample, which was run only once). Averages in gra boxes are corrected to full-strength.

Using the quantification standards established in Experiment 1, we can estimate infection intensity from zooplankton tows. Pachena Lake had an infection burden of 2.55 x 10^-4^ (31.9 copepods, 0.00813 *S. solidus*) and Black Lake had an infection burden of 7.03 x 10^-3^ (60.2 copepods, 0.0423 *S. solidus*). Multiplexed ddPCR analysis of eDNA from filtered lake water (Blackwater Lake, Vancouver Island B.C.) also showed positive signals for both host and parasite DNA. The eDNA sample was run only once and undiluted. ddPCR detected 0.42 copies of *S. solidus* DNA/μL and 1,767 copies/μL of cyclopoid host DNA in the environment.

## Discussion

Despite enormous control efforts, helminth parasites continue to infect billions of humans and animals annually (Lustigman et al. 2012, WHO 2017). One of the leading factors responsible for ineffective management is the failure to monitor parasites in their initial hosts (Webster et al. 2015). This gap in knowledge is central to understanding the factors that modulate the establishment and transmission of helminths. In this work, we designed and applied a ddPCR approach to detect and quantify host and parasite loads from environmental samples. We used a zooplankton-tapeworm model system as a case study to demonstrate the application of ddPCR as a valuable epidemiological tool. Small zooplankton (copepods, *A. robustus, ∼*1.3 mm) are the initial hosts to the larval stages of the helminth tapeworm *S. solidus.* Our assay can reliably detect infection in initial hosts starting from 0.1 picograms of total DNA. The assay is robust, demonstrating successful detection of both host and parasite DNA in both aquatic field samples (multi-species zooplankton tows) and eDNA samples (water filters). Our multiplexed ddPCR approach unites a cutting-edge method that has yet to be widely applied in disease ecology or epidemiological contexts. Together, we provide a solid framework for environmental detection and tracking that could greatly bolster management efforts of NTDs globally (Brindley et al. 2009).

### A tool for population-wide assessment (of both hosts and parasites)

ddPCR enables the quantification of infection loads in initial hosts at both the individual- and population-levels. Such data are crucial for understanding, predicting, and managing disease outbreaks. However, previous tools have failed to provide the level of quantitative detail required for such analyses. In helminths, for example, previous epidemiological efforts estimated infection dynamics in natural populations by quantifying zooplankton abundance (e.g., zooplankton density, Stutz et al. 2014) or using visual screening of infected zooplankton (e.g., Rusinek et al. 1996, Hanzelová and Gerdeaux 2003) to estimate infection levels and transmission rates in primary hosts. While these data are fundamental for parameterizing transmission models, estimating infection using these methods is labor intensive and time consuming. Our method helps address these logistical challenges by providing a high-throughput and cost-effective toolkit that can not only rapidly assess infection levels across multiple scales of biological organization but also detect low levels of infection that are often missed by canonical methods.

Previous studies have established data for *S. solidus* infection in secondary hosts (stickleback fish: Marcogliese 1995, Fuess et al. 2021). Our study contributes a fundamental piece of the puzzle to fill key gaps relating to initial infection and transmission dynamics of *S. solidus* and has the capacity to do the same for related host-parasite systems. This work will be applicable to estimating copepod contributions to R_0_, as we can leverage ddPCR outputs to estimate infection prevalence data for the initial host (Fenton et al. 2015). For example, we observed notable variation in infection burdens across different months and sampling sites (PCH mean June = 2.55 x 10□□ vs. BLA mean March = 7.03 x 10□^3^). Infection burden calculations are based on estimates of host numbers, which we were able to derive from our infection standard converting gene copy numbers to animal estimates. Using this standard, we found host numbers to vary considerably between sites and time-points in natural populations (PCH mean June = 31.9 cyclopoid copepods vs. BLA mean March = 60.2 cyclopoid copepods). Ultimately, evolutionary processes occur at the population level; without understanding infection dynamics in initial host populations, we miss critical eco-evolutionary processes that drive the entire infection sequence (e.g., contact probability, dilution effect). This principle is relevant for all helminthiases and is a key barrier to successful management of many neglected tropical diseases.

### Applications to other helminth systems

Our goal was to establish a generalizable toolkit that can serve as a foundation for future research on helminths dependent on copepods as initial hosts. The primers and probe-primers we have designed here lay the groundwork for broader research in a diverse array of other helminth systems (Box 1). Our assay targets the 18S rRNA gene of both *S. solidus* and cyclopoid copepods, leveraging conserved sequences found across the animal kingdom. Since *S. solidus* is known to infect various cyclopoid species (Wedekind 1997), we designed the Cyclopoid primer to encompass ecologically relevant hosts, such as *Acanthocyclops* sp. and *Macrocyclops* sp. These primers and probe-primers can be immediately applied to other helminthic diseases involving cyclopoids, including guinea worms and gnathostomiasis (Box 1, Nithiuthai et al. 2004). Additionally, our parasite primers broadly amplify the phylum *Platyhelminthes*, making them suitable for detecting other cestode helminths, such as broad tapeworms. Given the projected increase in helminth infections due to climate change and land-use disturbance, it is crucial to quantify animal reservoirs of these parasites (Blum and Hotez 2018).

### Limitations

Previous work optimizing ddPCR similarly found assays to be reproducible and sensitive in detecting rare or cryptic symbionts (Yang et al. 2014, Hiillos et al. 2021). While our assay demonstrated a strong linearity across the dilution gradient for both targets, reliability of target quantification strongly decreased in lower concentrations of total DNA (i.e., < 1pg of total DNA, helminth gene estimates differed by an order of magnitude when corrected compared to less dilute samples). For this reason, when handling field samples with unknown host/parasite concentrations it is important to establish adequate dilution to ensure proper droplet separation while maximizing target detection. In a limnological context, we found that field samples diluted to the 1 and 0.1ng/μL of total DNA resulted in the best detection and most reliable gene quantification within the ddPCR framework.

It is worth mentioning that one trade-off of this assay compared to manual inspection of copepods is that we do not know the stage-structure nor the sex of detected Cyclopoids. If a genetic marker between males and females were known, then it could potentially be included as another element of ddPCR reaction. This could be achieved by using different probe concentrations in order to generate distinct amplitude peaks without having to add an additional fluorescent dye. As in other systems, infection levels in copepods vary significantly across males and females (females: Hanzelová and Gerdeaux 2003, males: Rusinek et al. 1996, Wedekind and Jakobsen 1998). Based on laboratory studies, male *A. robustus* appear more susceptible to infection than females (IS, CAF, personal observations) but the role of infection on downstream effects, such as reproduction and sex determination remains unknown.

### Future applications of ddPCR

Helminthiases are notoriously difficult to track in the environment. Due to the relative rarity of helminth infections in the wild (Marcogliese 1995), a multiplexed ddPCR design quantifying both host and parasite prevalence allows for the reduction of false positives (e.g., non-ingested helminths) and the quantification of infection intensity in a sample. We found that multiplexing samples does not impede rare target detection. We use the multiplexed approach by using two dyes (HEX and FAM) to detect two targets; while most ddPCR instruments typically have only two fluorescence detection channels, it is possible to detect more targets by discriminating between the amplitude threshold of different target sequences. For example, we can discriminate between different, closely related helminth species (*S. solidus* and *S. cotti*) based on variations in fluorescence amplitude, even though both species are labeled with FAM dye (Supplementary File A, Fig. 2). An additional advantage of targeting the 18S rRNA gene is that it is present in multiple copies within an organism, which enhances detection sensitivity, particularly when the source is present in low abundance. This is especially valuable to detect infection in natural populations, as published infection burdens in copepods by other orders of cestodes are low (0.13-0.21 percent; Rusinek et al. 1996, Hanzelová and Gerdeaux 2003), and using our standard we find this intensity to be even lower in the *S. solidus* system (0.0002 and 0.007 percent).

We show here ddPCR is a powerful detection tool across diverse environmental samples. ddPCR offers applications of broad interest to ecologists beyond this scope, such as quantifying gene expression. This is especially applicable in helminth systems, as there has been significant uptick in the analysis of functional genomics in these worms that infect over 20 percent of the world’s population (Jolly et al. 2007). For instance, measuring growth-related gene expression in helminths (such as TRIP12 in *Schistosoma*, Gobert et al. 2006) could indicate their readiness for their transmission to the next host, providing *in situ* insights on parasite life history and transmission dynamics. Additional genome sequencing and gene expression studies of initial hosts will also set the foundation for many studies relevant to population ecologists. For example, identifying growth-markers in copepods (e.g., pre vs. post metamorphosis) would also open up avenues to adding stage-structured analysis to field samples, allowing researchers to consider population demographics of diverse hosts in natural populations.

### Conclusions

This study demonstrates multiplexed ddPCR as a highly sensitive and repeatable method to simultaneously quantify parasite and host genes from multi-species, environmental samples. By establishing a standard that translates gene copy numbers into actual animal estimates, our assay offers a valuable tool for informing management decisions across helminth systems. We present a toolkit of primers and probes that are applicable to a range of helminths species, offering a flexible toolkit for studying NTDs and host-parasite interactions in natural systems. Future work may build upon this methodology by considering additional target species (i.e., co-infection) by varying probe concentrations in addition to multiplexing assays. By bridging molecular precision with ecological (and societal) relevance, this study contributes to promoting the early detection and quantification of helminthiases globally.

## Acknowledgements

This work was funded in part by National Science Foundation EEID (Grant Number 144- 873100-4-AAL3614, AH, DIB, JLH) and the University of Wisconsin, Madison (101-870663-4, startup funds to JLH). We are thankful to the Carleton College Career Center Summer Internship Funding for additional support provided to JS. We would like to thank Carol Enumi Lee and her UW lab for providing us with calanoid copepods. Copepod icons throughout the paper were modified from original design by BioRender. Lastly, this manuscript would not have been possible without the music of *Wings*, which helped CAF and ENE persevere through many troubleshooting steps.

## Author Contributions

CAF wrote the initial manuscript, all authors contributed to manuscript revisions and approved of the final version of the submission. CAF and ENE designed molecular components of ddPCR assay and performed molecular analysis. CAF designed the figures. IS performed infection assays and lead generation of infection standards, supported by JS and JC. CW and JW contributed helminth tissues and live cyclopoid copepods and cestodes. AH, JBIII, HA, EC, DIB, and JLH contributed field samples (eDNA and zooplankton tows).

## Data Accessibility

Raw ddPCR data and R code are available in the Environmental Data Initiative (EDI) repository (DOI TBD) and CAF’s GitHub (https://github.com/chloefouilloux/18s_ddPCR) for ease of access. All data is publicly available, ensuring that researchers can access, reuse, and reproduce the analyses.

## Benefit sharing

Benefits generated: The research addresses a priority concern of identifying tapeworm helminths of global health concern in natural habitats. All data will be shared with the broader public via appropriate biological databases. All undergraduate collaborators of this manuscript are included as co-authors, supporting the early career of our next generation of scientists.

## Notes

https://portal.edirepository.org/nis/mapbrowse?scope=edi&identifier=1855&revision=1

### Competing Interest Statement

The authors have declared no competing interest.

