## Supplementary File C for "Needle in a haystack: A droplet digital polymerase chain reaction assay to detect rare helminth parasites infecting natural host populations"

**Supplementary File C. Total *Schistocephalus solidus* loads (both encysted and free) within threespine stickleback.**

In Fouilloux et al. (2025) authors design both probe and probe-primers for an assay to detect *Schistocephalus solidus* parasites. Here, authors justify the lakes highlighted in the main text as a result of the *S. solidus* infection trends in fish (although sampled at different time points), the secondary obligate host of *S. solidus.*

**Tapeworm quantification.** *S. solidus* were counted from dissected three-spine sticklebacks. Here, we include both free and encysted tapeworms within fish.

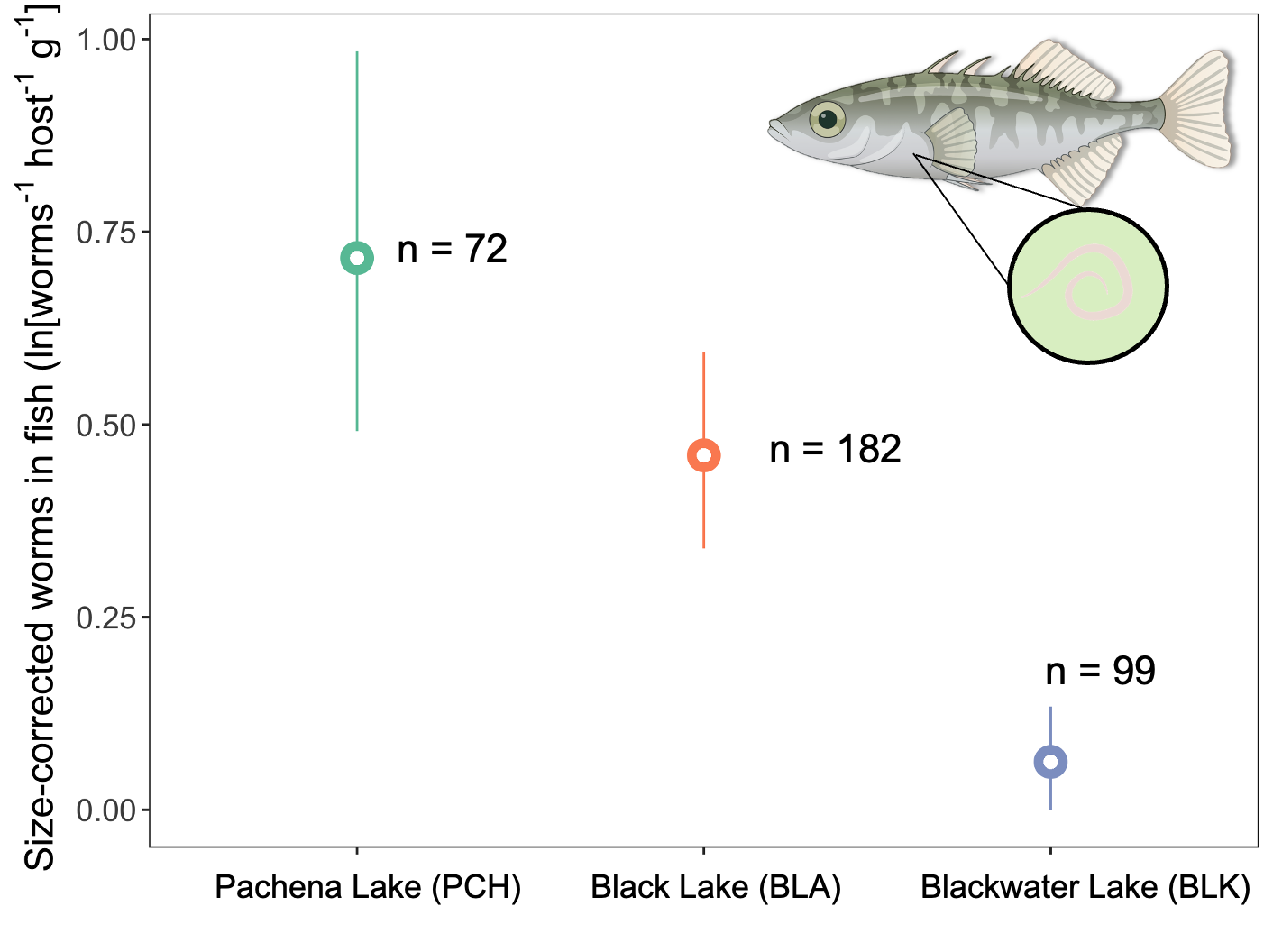

**Fig 1. Size-corrected *S. solidus* loads in fish.** Point ranges indicate lake-level means with 95 CIs. Sample sizes indicate the number of fish contributing to lake-level mean.
