## Supplementary File B for "Needle in a haystack: A droplet digital polymerase chain reaction assay to detect rare helminth parasites infecting natural host populations"

**Supplementary File B. Specificity of cyclopoid primers to other genera of copepods.**

In Fouilloux et al. (2025) authors design both probe and probe-primers for an assay to detect Cyclopoid copepod hosts. Here, authors empirically test primer and probe-primer amplification of Cyclopoids (*Acanthocypolds robustus*) versus Calanoids (*Eurytemora affinis*).

**TL;DR** PCR probes do not discriminate between genera but ddPCR probe-primers do.

**DNA Extraction**

Same method as detailed in the main text was used for a *E. affinis* standard containing 100 individuals.

**PCR primer testing**

Targeting the 18S rRNA gene, the designed **primer set** (517F forward, C138R) more strongly amplifies cyclopoid template over calanoids (Fig 1). However, there is a 154-nucleotide fragment that appears (although to a much lesser extent) in calanoids, preventing discrimination of the two genera using only PCR primers.

**PCR Reaction details**

PCR reactions were performed in a total volume of 25μL [12.5μL Taq Master Mix (PR1MA PR1001-R, MidSci), 8.5μL PCR water, 1.0μL each forward and reverse primer (10 μM conc.), 2μL DNA (10ng μL^-1^ conc.) (or 10.5μL PCR water for NTC)] in a BioRadT100 thermal cycler. PCR conditions included a denaturation step of 3 min at 95°C, followed by annealing and elongation step of 40 cycles at 95°C for 15s, 55°C for 15s, 72°C for 30s, and a final step of 72°C for 5 min. PCR products were run on 1.6% agarose gel using SYBR safe staining (Invitrogen). During optimization primer sets were tested across a temperature gradient (55-62.5C) of annealing temperatures, but the 154-nt band persisted in calanoid samples across different cycling conditions.

**ddPCR probe-primer testing**

The designed probe-primer (**HEX**-CAGTCGCGTCAAATAACGGACCGCCC-**IBFQ**) is specific to cyclopoids and excludes calanoids (Fig 2). ddPCR assay conditions are identical to those detailed in the main text. For experimental purposes, annealing was run at a gradient spanning 62.5, 60, and 57.5C. There was no detection of calanoid DNA at any temperature.

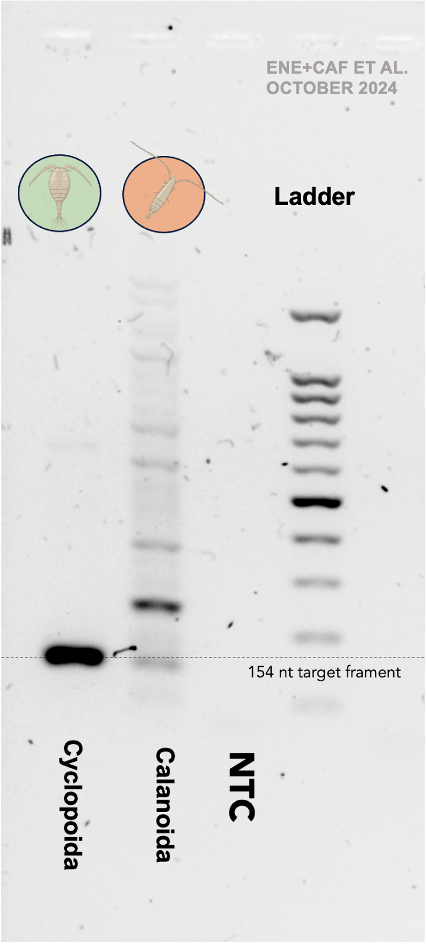

**Fig 1. PCR gel of Fouilloux et al. 2024 primers on cyclopoid vs. calanoid DNA.** Light 154 nucleotide band in calanoid lane prevents genera to be discriminated using only PCR primers.

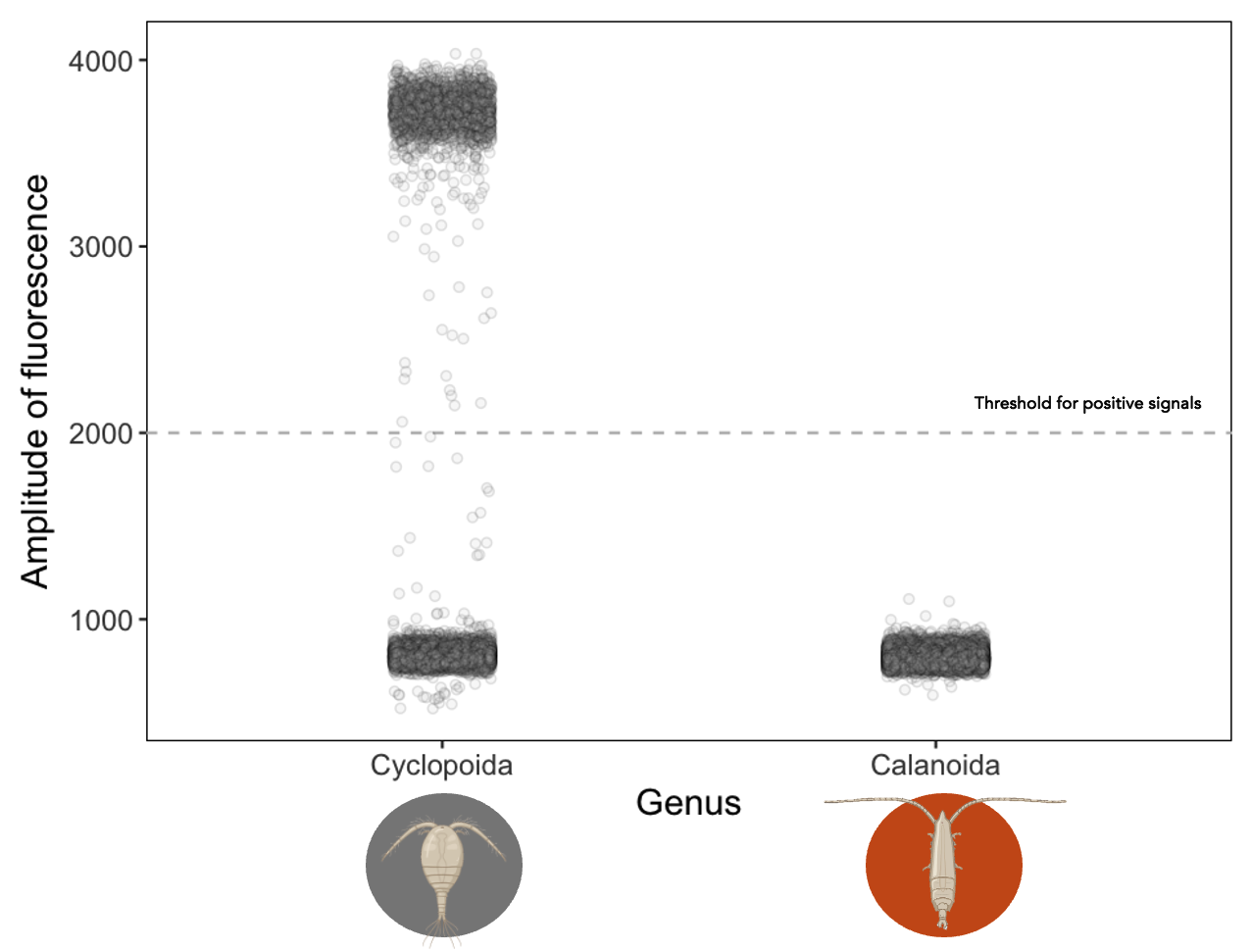

**Fig 2. ddPCR raw amplification plot of a single cyclopoid and calanoid well.** Probe-primers did not amplify calanoid DNA, allowing for discrimination of genera.
