## Supplementary File A for "Needle in a haystack: A droplet digital polymerase chain reaction assay to detect rare helminth parasites infecting natural host populations"

**Supplementary File A. Specificity of *Schistocephalus solidus* primers to other species of *Schistocephalus* tapeworms.**

In Fouilloux et al. (2025) authors design both probe and probe-primers for an assay to detect *Schistocephalus solidus* parasites. Here, authors empirically test primer and probe-primer amplification of three *Schistocephalus* species: *S. solidus, S. cotti, and S. pungitii.*

**TL;DR** PCR probes do not discriminate between tapeworm species. ddPCR probe-primers can discriminate between *S. solidus* and *S. cotti* but not between *S. solidus* and *S. pungitii.*

**DNA Extraction**

DNA from *S. cotti* and *S. pungitii* came from tissue samples stored in 90% EtOH. DNA was extracted using a Qiagen DNeasy Blood and Tissue Kit following kit instructions.

**PCR primer testing**

Targeting the 18S rRNA gene, the designed **primer set** (F517 forward, S708R) amplifies all 3 *Schistocephalus* species (Fig 1). To check cross-species specificity, we included some Cyclpoid copepod DNA. There was some nonspecific binding of copepod DNA, but none of it overlapped with the target fragment size for *Schistocephalus.*

**PCR Reaction details**

PCR reactions were performed in a total volume of 25μL [12.5μL Master Mix, 8μL PCR water, 12.5μL forward and reverse primer (10 μmol l^-1^ conc.), 2μL DNA (10ng μL^-1^ conc.) (or 8.5μL of PCR water for NTC)] in a BioRadT100 thermal cycler. PCR conditions included a denaturation step of 5 min at 95°C, followed by annealing and elongation step of 40 cycles at 95°C for 45s, 54°C for 45s, 72°C for 60s, and a final step of 72°C for 7 min with a 1°C/sec ramp speed. PCR produced were run on 1.5% agarose gel using SYBR safe staining (Invitrogen).

**ddPCR probe-primer testing**

The designed probe-primer (**FAM-**CCCACCTACACCGACACAATCAGCC**-IBFQ**) amplifies all three *Schistocephalus* species but at different efficiencies (Fig 2). ddPCR assay conditions are identical to those detailed in the main text. For experimental purposes, annealing was run at a gradient spanning 62.5, 60, and 57.5C. Here, *S. cotti* can be distinguished from *S. solidus* and *S. pungitii.* While *S. solidus* and *S. pungitii* cannot be distinguished genetically, the two species have never been observed to co-occur on Vancouver Island (DIB personal observation).

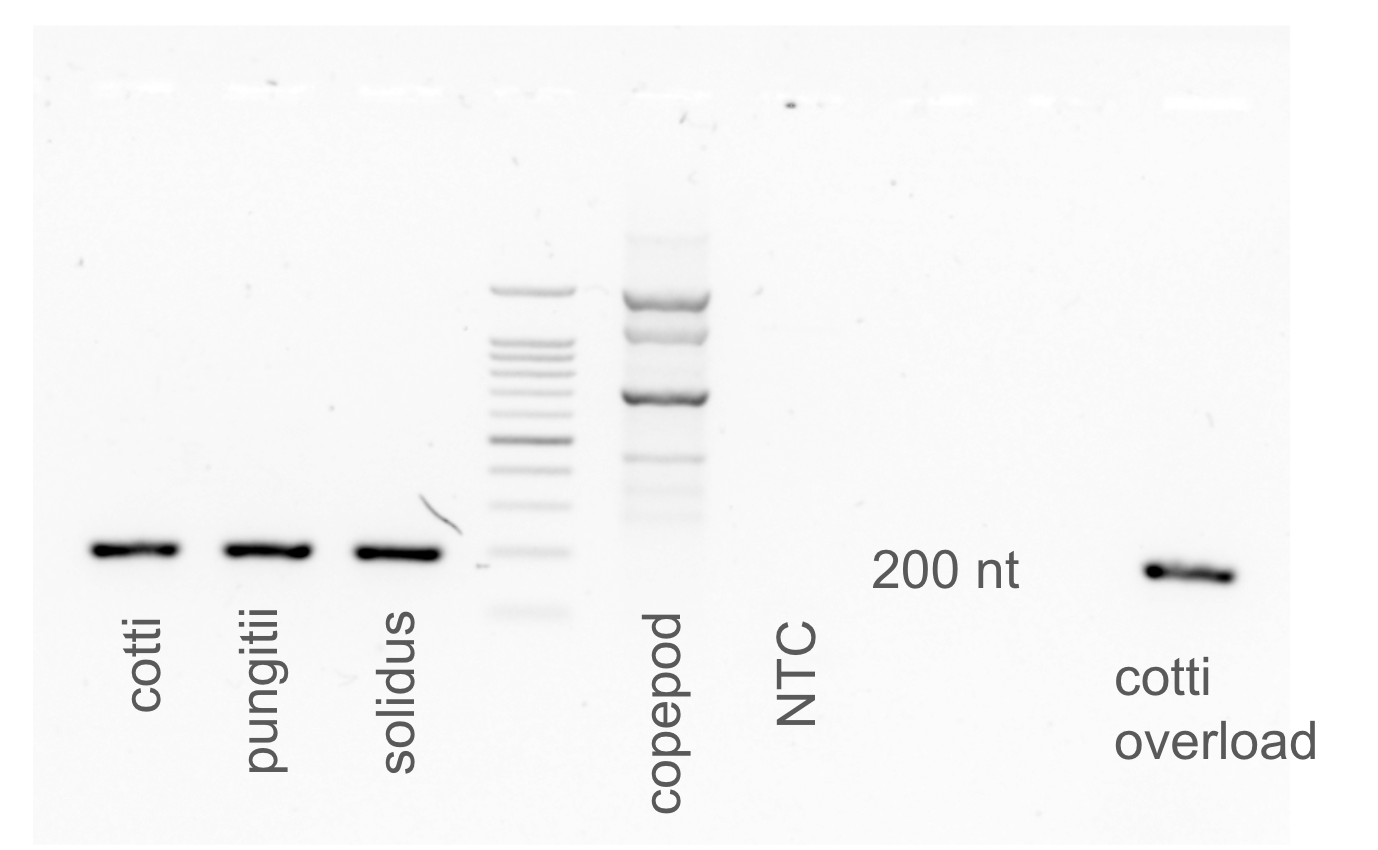

**Fig 1. PCR gel of Fouilloux et al. 2024 primers on diverse *Schistocephalus sp*.** PCR primers are equally suitable to amplify all three helminth species.

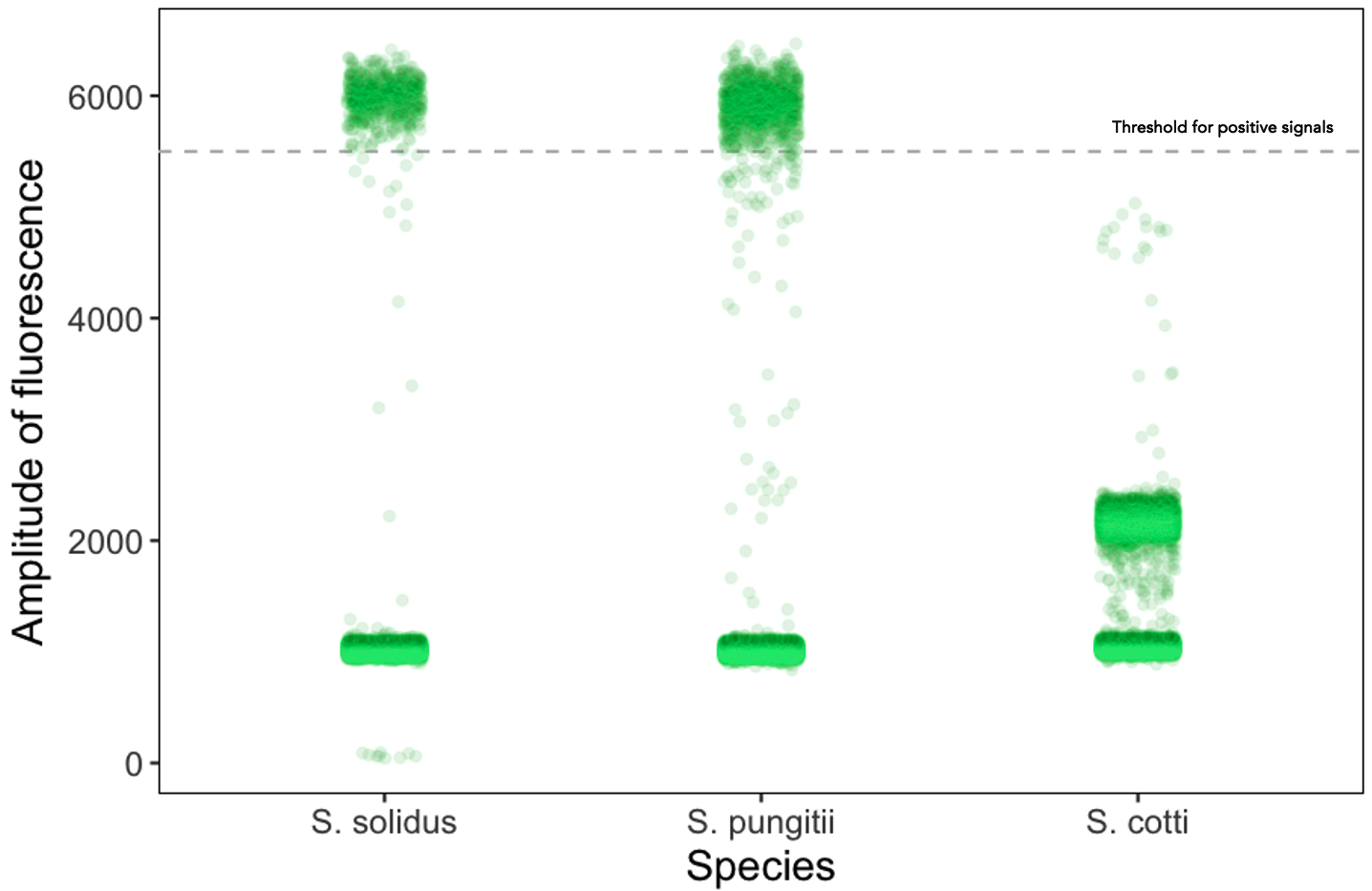

**Fig 2. ddPCR raw amplification plot of *S. solidus, puntii,* and *cotti* selected wells.** Probe-primers amplify all of the helminth DNA, however, the amplitude of fluorescence between the species allows for the discrimination of *S. cotti.* Here can discriminate between species labeled with the same dye (FAM) by differences in the amplitude of fluorescence.
